# Metabolic Rewiring by α-Synuclein Enables Mitohormetic Protection

**DOI:** 10.64898/2025.12.05.692538

**Authors:** Shay Geula, Inna Grosheva, Yehudit Zaltsman, Limor Regev, Sabita Chourasia, Atan Gross

**Affiliations:** Department of Immunology and Regenerative Biology, Weizmann Institute of Science, Rehovot, Israel 76100

**Author notes:** Co-first authors (Shay Geula, Inna Grosheva, Yehudit Zaltsman). Corresponding author: Atan Gross, PhD Department of Immunology and Regenerative Biology Weizmann Institute of Science Rehovot, Israel 76100.

## Abstract

α-Synuclein (αSyn) is widely associated with Parkinson’s disease pathology, yet its physiological impact on cellular metabolism remains unclear. Here we show that stable αSyn expression in HEK293 cells induces coordinated metabolic remodeling that enhances mitochondrial resilience. αSyn interacts with lactate dehydrogenase A (LDHA) and αSyn–expressing clones exhibit elevated LDHA activity. Moreover, these clones exhibit increased lactate secretion, enhanced glycolysis, and reduced mitochondrial-reactive oxygen species, coupling metabolic rewiring to improved mitochondrial adaptation. Strikingly, following chronic low-dose rotenone preconditioning, αSyn– expressing clones acquire robust resistance to subsequent respiratory complex I inhibition, revealing a potent αSyn–dependent mitohormetic response. These findings identify αSyn as a conditional metabolic modulator that supports mitochondrial adaptation under sustained stress.

## Introduction

Mitochondrial dysfunction and oxidative stress are major cellular disturbances associated with the onset and progression of Parkinson’s disease (PD) [1]. Disruption of mitochondrial complex I specifically in dopaminergic (DA) neurons induces progressive parkinsonism in mice [2], and environmental inhibitors of oxidative phosphorylation (OXPHOS) recapitulate PD-like pathology [3, 4], establishing mitochondrial impairment as a driver of PD development. PD is also characterized by the presence of Lewy bodies, whose major protein component is α-synuclein (αSyn)[5]. Mutations or multiplications of the SNCA gene cause autosomal-dominant PD, demonstrating that increased or altered αSyn expression is sufficient to initiate disease [6].

Misfolded or aggregated αSyn exerts toxic gain-of-function effects: αSyn preformed fibrils (PFFs) seed aggregation, recruit endogenous αSyn into Lewy-like inclusions, and trigger synaptic dysfunction and neurodegeneration [7, 8]. Aggregated αSyn can also impair mitochondrial respiration by inhibiting complex I activity [9]. Importantly, however, monomeric αSyn alone is not sufficient to produce PD-like pathology in-vivo. In a recent study, mice injected with monomeric αSyn into the striatum did not lose DA neurons and did not develop motor impairments, despite the presence of αSyn protein, supporting the conclusion that monomeric αSyn by itself is not neurotoxic [10].

Despite αSyn’s central involvement in PD pathology, extensive evidence shows that soluble, physiological αSyn is essential for normal neuronal function and can exert protective roles under stress. αSyn is highly enriched at presynaptic terminals, where it binds synaptic vesicle membranes, regulates vesicle clustering, and promotes SNARE-complex assembly via interaction with VAMP2/synaptobrevin-2 [11]. Mice lacking α-, β-, and γ-synuclein exhibit synaptic dysfunction, reduced SNARE-complex formation, and age-dependent neurodegeneration, indicating that synucleins are required for long-term neuronal integrity [12]. Several experimental studies demonstrate that αSyn is cytoprotective under oxidative stress: overexpression of αSyn inhibits the c-Jun N-terminal kinase (JNK) pro-apoptotic pathway through induction of the scaffold protein JIP-1b [13]; chronic oxidative stress upregulates αSyn expression, correlating with reduced neuronal degeneration [14].

DA neurons are among the most metabolically demanding neurons in the brain. They rely heavily on glucose as their primary energy source, making glucose metabolism critical for their survival and function. Glucose metabolism is impaired in PD [15–18], and metabolic shifts from OXPHOS to glycolysis have been shown to occur in neurons and to exert protective effects. For example, aerobic glycolysis in neuronal somata has been shown to reduce oxidative damage [19], glycolytic preconditioning in astrocytes mitigates trauma-induced neurodegeneration [20], and enhanced glycolysis through activation of phosphoglycerate kinase 1 (PGK1) attenuates PD progression in multiple models [21]. Furthermore, αSyn itself has been shown to modulate glycolysis: it induces microglial migration via PKM2-dependent glycolysis, directly enhancing glycolytic flux [22]. In adipocytes, αSyn stimulates glucose uptake and utilization through activation of the Gab1/PI3K/Akt pathway [23], demonstrating that αSyn can modulate upstream glucose-signaling pathways as well.

Hypoxia, which shifts metabolism from OXPHOS toward glycolysis, provides another paradigm whereby metabolic rewiring protects neurons [24, 25]. In addition, hypoxia was recently shown to ameliorate neurodegeneration and motor impairment in mice injected with aggregated αSyn, despite the persistence of αSyn aggregates[10]. These findings indicate that hypoxia alleviates PD pathology upstream of αSyn aggregation, possibly by reducing the toxic burden of oxygen associated with mitochondrial dysfunction. Hypoxia stabilizes hypoxia-inducible factor 1α (HIF-1α), which induces transcriptional reprogramming toward glycolysis [26]. A key HIF-1α target is lactate dehydrogenase A (LDHA) [27], which converts pyruvate to lactate and regenerates NAD⁺ under low-oxygen conditions, thereby sustaining glycolytic ATP production.

Here, we demonstrate that αSyn interacts with LDHA, and that αSyn-expressing clones show elevated LDHA activity, increased lactate secretion, and reduced mitochondrial reactive oxygen species, both prior and post treatment with rotenone, a respiratory complex I inhibitor. Moreover, αSyn-expressing clones exhibit higher glycolytic and mitochondrial respiratory activity and have a proliferative advantage over control cells following preconditioning with low dose rotenone. Thus, αSyn enhances both glycolytic and oxidative metabolic pathways, at least in part via LDHA stimulation, providing cells with increased energetic flexibility during mitochondrial stress.

## Results

### αSyn directly interacts with LDHA

To study the cellular biology of αSyn in cells, we generated HEK293T stable clones expressing αSyn or carrying an empty vector. Cells were stably transduced with αSynIRES-GFP or GFP lentivirus, FACS sorted to isolate cells with high GFP expression levels, and single-cell clones were established for long term culture. αSyn expression was detected via Western blot and immunofluorescent labeling with anti-αSyn Ab (Supp Fig 1A, B). Importantly, the GFP signal correlated with αSyn labeling (Supp Fig 1B), making GFP intensity a useful readout to monitor αSyn expression. αSyn stable overexpression did not affect cell growth, and expression of the transgene remained constant upon cell culturing in regular conditions (data not shown).

To assess the involvement of mitochondrial stress in triggering αSyn aggregation, we exposed αSyn-expressing clones to rotenone, a respiratory complex I inhibitor. Three αSyn-expressing clones (C1-C3) were either treated with DMSO or with 10µM rotenone for 4hrs, harvested, and treated with the disuccinimidylglutarate (DSG) non-reversible protein cross-linker. We used the DSG cross-linker since a previous report used it to successfully identify oligomeric forms of αSyn [28]. The cells were then lysed, and Western blot analyzed using anti-αSyn Abs. In cells treated with DSG, we detected several highmolecular-weight (HMW) αSyn-immuno-reactive bands, which in two of the three clones were enhanced following rotenone treatment (Fig 1A). These HMW bands might represent αSyn homo-oligomers and/or hetero-oligomers between αSyn and another protein(s).

To gain further evidence that p17-αSyn was indeed part of these HMW-crosslinked bands/complexes, we isolated the cytosolic S100 fraction from αSyn-clone #2, treated it with DSG and collected samples at time 0, 10, 15, 20, 30 min. This time-course experiment showed a gradual decrease in the intensity of the p17-αSyn-band concomitantly with the appearance/increase in the intensity of the HMW αSyn-immunoreactive bands (Fig 1B). These results provide further evidence that the HMW-cross-linked bands/complexes contain p17-αSyn. These results also suggest that part of the HMWcross-linked bands/complexes detected in whole cells are comprised of cytosolic protein(s).

**Figure 1.**
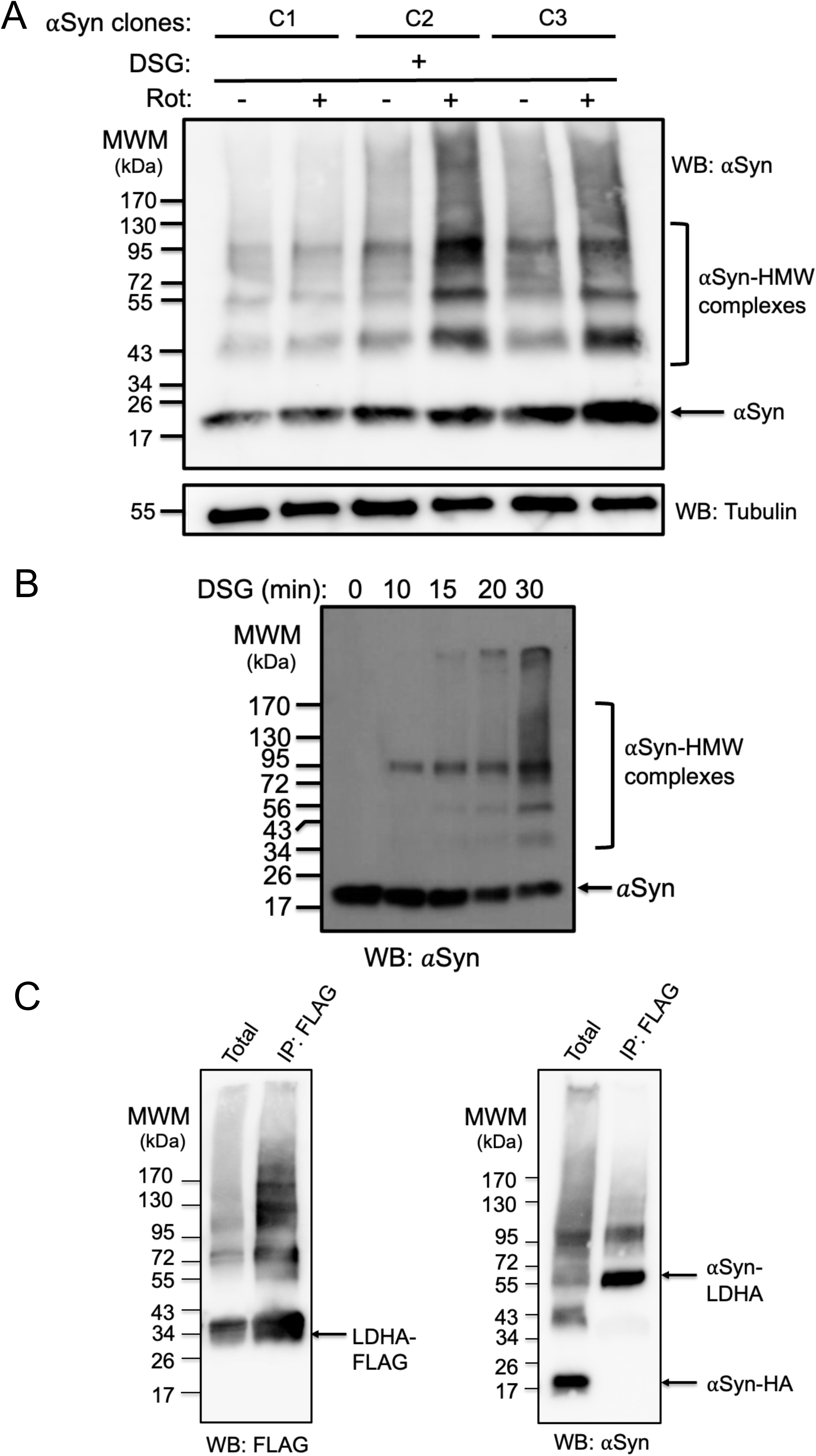
αSyn directly interacts with LDHA. (A) Three HEK293T αSyn-expressing clones were treated with either DMSO (-) or with 10 µM rotenone (+) for 4 hrs, cross-linked with DSG, and analyzed by Western blot using anti-αSyn antibody. Under basal conditions several cross-linked high-molecular-weight (HMW) αSyn-reactive species were detected, which increased following rotenone treatment. (B) Time-course DSG cross-linking of cytosolic S100 fraction isolated from αSyn C2 (0– 30 min) showed a progressive loss of the intensity of the p17-αSyn-band concomitantly with the appearance/increase in the intensity of the HMW αSyn-immuno-reactive bands. (C) HEK293T cells were co-transfected with αSyn-HA and LDHA-Flag, cross-linked with DSG, pull down using anti-Flag Abs, eluted with Flag peptide, and analyzed by Western blot with either anti-Flag Abs (left panel) or anti-αSyn Abs (right panel). Total cell lysate after cross-linking is shown in lane 1; lane 2 shows anti-Flag Ab immunoprecipitants. αSyn-HA cross-linked to LDHA-Flag appears in a ∼50kDa complex (right panel).

To identify the protein composition of the HMW αSyn-immuno-reactive bands detected in whole-cell lysates, we scaled up their detection by generating a C-terminal HA-tagged αSyn construct, transiently expressing it in naïve HEK293T cultured cells, treating the cells with the DSG cross-linker, and affinity-purifying HA-tagged αSyn using an anti-HA antibody matrix. Bound proteins were subsequently eluted with HA peptide, separated by SDS–PAGE, and the gel was stained with InstantBlue® Coomassie (Supp Fig 1C). The four detected/indicated bands were then cut out from the gel and processed for mass spectrometry analysis.

As expected, mass spectrometry analysis showed that all four bands (especially the ∼20kDa band) were enriched for αSyn peptides (see Table 1). Also heat shock proteins 70 and 90 were ranked high. Interestingly, several glycolytic enzymes – Alpha-enolase, Pyruvate kinase, Fructose-bisphosphate aldolase A, Lactate dehydrogenase A and B chains, Glyceraldehyde-3-phosphate dehydrogenase, and Glucose-6-phosphate isomerase - were also ranked high. Our earlier results, showing that the intensity of the HMW αSyn-immuno-reactive bands increases in response to rotenone, and the known fact that treatment with rotenone results in a compensatory increase in glycolysis [29], suggested that monomeric αSyn is possibly involved in modulating glycolysis by binding/activating one or more of these glycolytic enzymes.

Notably, a key enzyme that enables continuous high levels of glycolysis during mitochondrial stress, such as hypoxia, is Lactate dehydrogenase A (LDHA), that converts pyruvate to lactate [30]. Under hypoxia conditions, hypoxia-induced factor 1α (HIF-1α) is stabilized and one of its key transcriptional targets is LDHA [27]. Moreover, peptides of 36kDa-LDHA were enriched in the mass spec analysis of part of the cross-linked bands/complexes (Table 1). Based on all the above, we decided to further investigate the interaction between αSyn and LDHA.

To assess whether αSyn directly binds to LDHA, cells were co-transfected with αSyn-HA and LDHA-Flag plasmids, treated with DSG, lysed and co-immunoprecipitated with anti-Flag Abs followed by Western blot using either anti-Flag or anti-αSyn Abs. As expected, pull-down of LDHA-Flag with anti-Flag antibodies resulted in a marked enrichment of high-molecular-weight LDHA complexes (Fig 1C, left panel). Interestingly, and in agreement with the mass spectrometry results, αSyn-HA was also detected in the LDHA-Flag pull-down as part of a ∼50kDa complex (Fig 1C, right panel), indicating that p17-αSyn directly associates with p36-LDHA.

### Following chronic low-dose rotenone preconditioning, αSyn–expressing cells acquire robust resistance to subsequent mitochondrial inhibition

The results presented above suggesting that monomeric αSyn is possibly involved in increasing glycolysis by binding LDHA, prompted us to assess whether the αSyn-expressing clones are less sensitive than the vector-expressing clones to rotenone.

We observed no differences in the rates of cell death between the groups when treated with either 1µM or 100nM rotenone. Thus, next we assessed the effect of a nonlethal dose of 40nM rotenone. Two weeks treatment of vector- and αSyn-expressing clones with 40nM rotenone resulted in enrichment of strong-GFP expressing cells only in cells expressing αSyn, as shown by fluorescent microscopy and by FACS analysis (Fig 2A, B, respectively). As expected from the GFP results, total αSyn levels were increased in cells continuously treated with 40nM rotenone, as detected by Western blot (Fig 2C).

**Figure 2:**
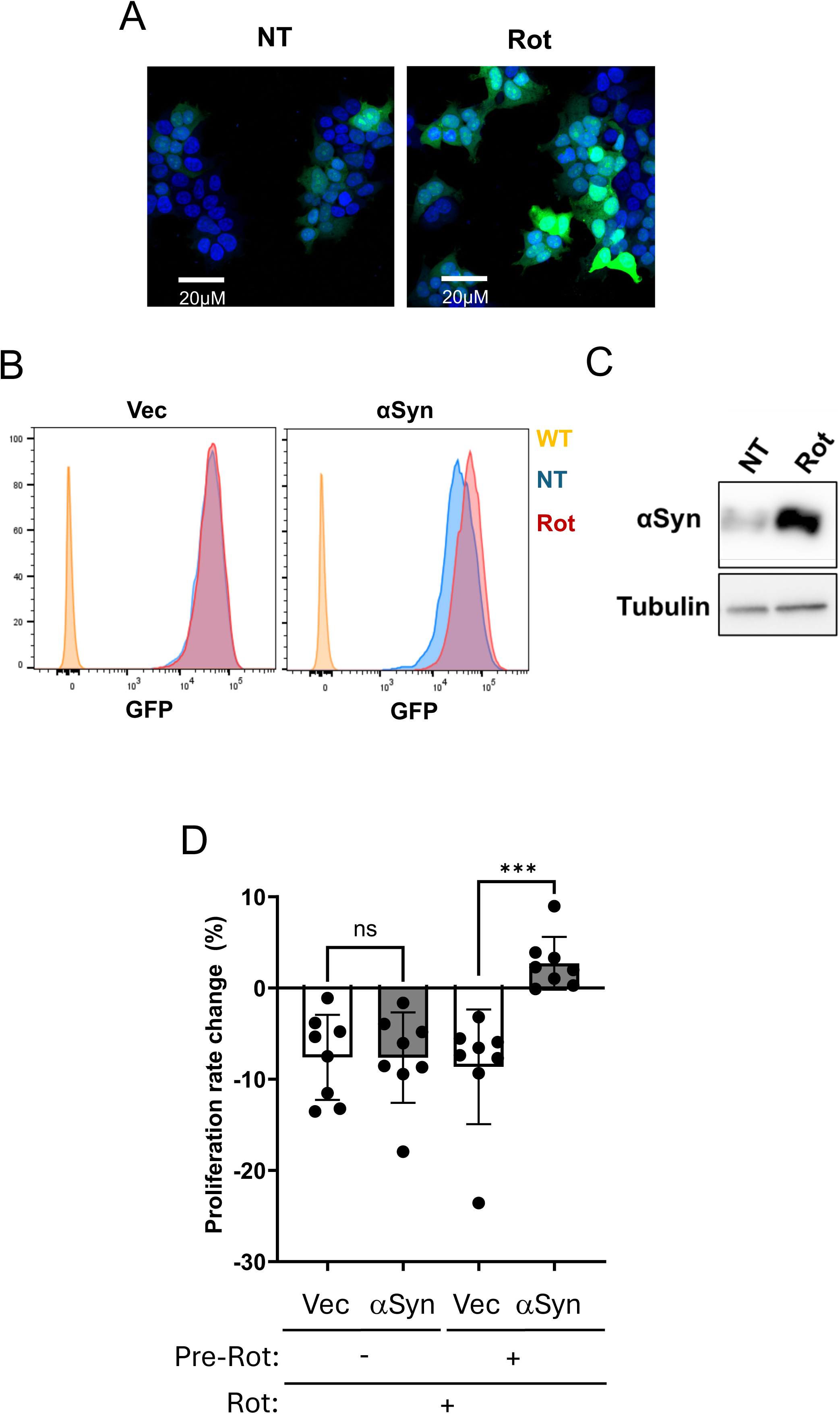
Following chronic low-dose rotenone preconditioning, αSyn–expressing cells acquire robust resistance to subsequent mitochondrial inhibition. (A) Fluorescence microscopy of αSyn-expressing clones in basal condition (NT) and after exposed to 40 nM rotenone for two weeks (Rot). Chronic rotenone treatment led to enrichment of high-GFP–expressing cells, indicating selective survival or expansion of αSyn-expressing cells. (B) Flow cytometry analysis of GFP intensity in vector- and αSyn-expressing clones under basal conditions (NT) and after two weeks of 40 nM rotenone treatment (Rot). Rotenone induces a shift toward higher GFP intensity specifically in αSyn-expressing clones. WT represents naïve wild-type HEK293T cells. (C) Western blot analysis of αSyn protein levels in vector- and αSyn-expressing clones before (NT) and after two weeks of 40 nM rotenone exposure (Rot). Rotenone increased αSyn levels in αSyn-expressing clones, consistent with the enrichment of GFP⁺/αSyn⁺ cells observed in A and B above. (D) Growth analysis of vector- and αSyn-expressing clones. A total of 2,000 cells were seeded into 96-well plates and monitored for four days using Incucyte® Live-Cell instrument. Prior to imaging cells were either cultured in basal media (Pre-Rot: -) or preconditioned with 40 nM rotenone for two weeks (Pre-Rot: +). Cell growth was monitored for 96 hours either in regular medium or in medium containing 40 nM rotenone. Proliferation rate for each condition was calculated for exponential growth phase. Rotenone induced a 5–15% reduction in proliferation rate in both groups upon acute treatment in the absence of preconditioning; however, growth of αSyn-expressing clones did not slow down in the presence of rotenone following metabolic preconditioning, indicating enhanced adaptation to complex I inhibition. Statistical significance: ns, not significant, ***P < 0.005.

These results suggested that high-αSyn-expressing cells are less sensitive to lowdose-rotenone treatment and therefore proliferate faster than low-αSyn-expressing cells. Next, we monitored cell proliferation in standard growth media and in media containing 40nM rotenone. As expected, rotenone slowed proliferation in both vector- and αSyn-expressing clones (5-15% decrease from growth in standard growth media), however there was no difference in the proliferation rate between the clones (Fig 2D, 1^st^ and 2^nd^ bar graphs).

Next, we assessed whether metabolic preconditioning with rotenone (Pre-Rot) would differentially affect the clones. Chronic exposure to 40nM rotenone for two weeks prior to the rotenone challenge had no effect on vector-expressing clones, which again showed a 5–15% decrease in proliferation rate (Fig 2D, 3^rd^ bar graph). Strikingly, αSyn-expressing clones preconditioned with 40nM rotenone for two weeks became resistant to rotenone-induced growth inhibition and even showed an increase in growth (Fig 2D, 4^th^ bar graph).

Taking together, these findings reveal a potent αSyn–dependent mitohormetic response, in which αSyn acts as a conditional metabolic modulator that supports mitochondrial adaptation under sustained stress.

### αSyn–expressing clones exhibit elevated LDHA activity, increased lactate secretion, and enhanced glycolysis

We next assessed whether LDHA plays a role in the αSyn–dependent mitohormetic response. To assess this point, we examined whether αSyn-expressing clones exhibit higher LDHA enzymatic activity, and higher levels of its product, lactate.

LDHA converts pyruvate to lactate, which inhibits pyruvate entry into mitochondria and sustains glycolysis by converting NADH to NAD^+^[31](Fig 3A, top left panel). We monitored LDHA activity by measuring the decrease in NADH (=OD 340nm). And indeed, all αSyn-expressing clones show faster consumption of NADH as compared to the vector-expressing clones, both prior and post two-weeks treatment with 40nM rotenone (Fig 3A, bottom left panels and right panel). These results are consistent with the idea that αSyn can activate a glycolytic rescue program even before overt mitochondrial dysfunction occurs.

**Figure 3:**
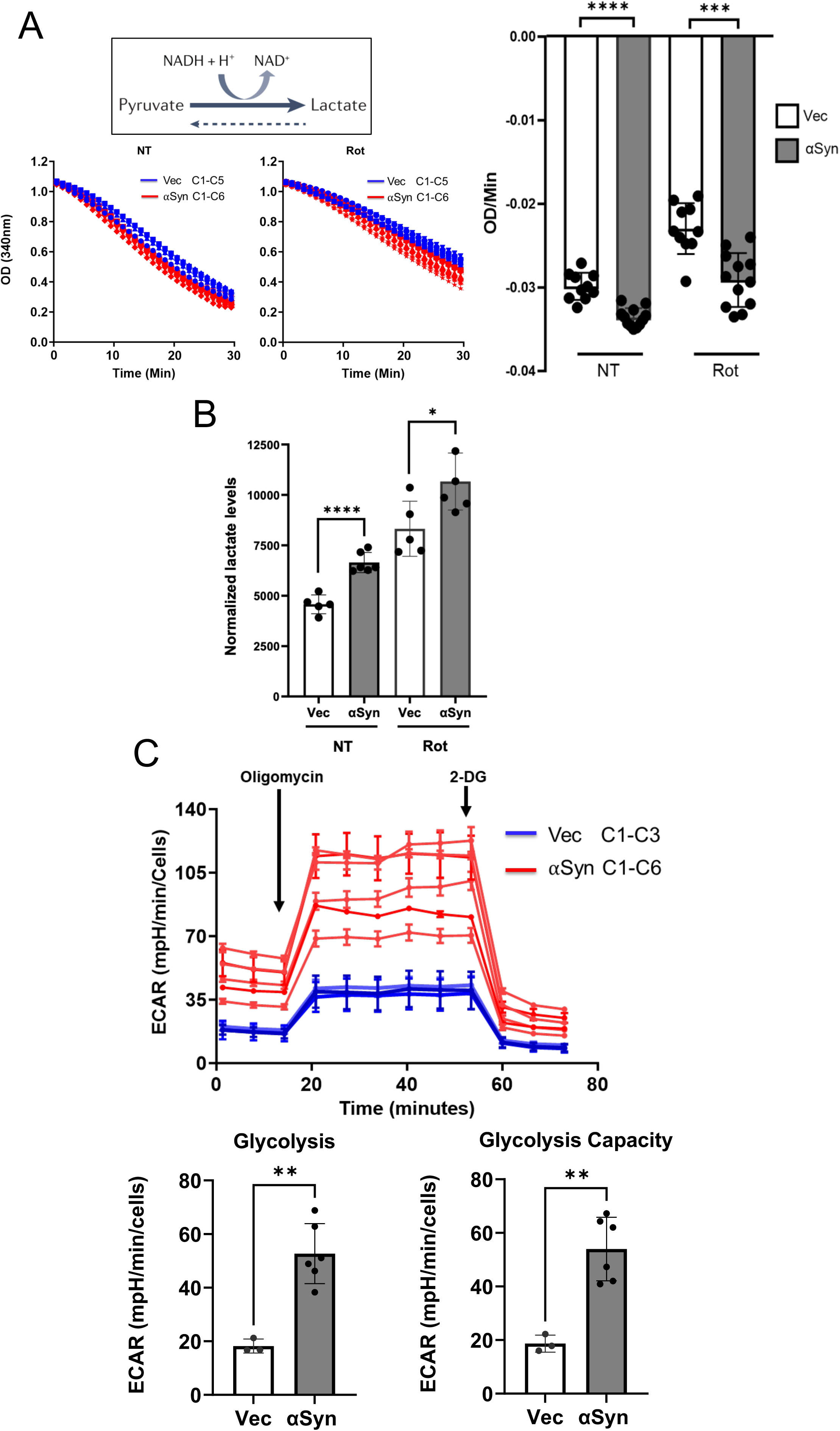
αSyn–expressing clones exhibit elevated LDHA activity, increased lactate secretion, and enhanced glycolysis. (A) *Top:* Schematic illustrating the LDHA-catalyzed reaction, in which LDHA converts pyruvate to lactate while oxidizing NADH to NAD⁺. *Bottom:* LDHA activity was assessed by monitoring the decrease in NADH absorbance at 340 nm. All αSyn-expressing clones showed faster NADH consumption as compared to vector-expressing clones under basal conditions (NT), indicating elevated LDHA activity. Two weeks of growth in the presence of 40 nM rotenone (Rot) reduced LDHA activity in both groups. *Right:* Quantification of LDHA activity in αSyn-expressing clones and vectorexpressing clones under basal conditions (NT) and after two weeks of growth in the presence of 40 nM rotenone (Rot). Each dot represents the slope of an individual reaction. Statistical significance: ***P<0.005, ****P<0.001. (B) Secreted lactate levels in conditioned medium were measured in vector- and αSynexpressing clones before (NT) and after two weeks of growth in the presence of 40 nM rotenone (Rot). Values were normalized to total protein content. αSyn-expressing clones exhibit higher lactate secretion under both prior and post rotenone treatment. Statistical significance: *P<0.05, ****P<0.001. (C) Seahorse ECAR analysis of vector- and αSyn-expressing clones. Basal glycolysis was measured following glucose addition, and maximal glycolytic capacity was determined after oligomycin injection. All αSyn-expressing clones displayed higher basal ECAR and higher glycolytic capacity compared to all vector-expressing clones. Statistical significance: **P<0.01.

LDHA converts pyruvate to lactate, and much of the lactate is secreted out of the cells [31]. Thus, we measured the levels of lactate in the cell medium, and as in the case with LDHA activity, all αSyn-expressing clones show higher levels of lactate in the medium as compared to the vector-expressing clones, both prior and post two-weeks treatment with 40nM rotenone (Fig 3B).

Next, we used the seahorse instrument to measure the levels of glycolysis (ECAR). As expected from the LDHA activity and lactate results, all αSyn-expressing clones show higher levels of glycolysis and glycolysis capacity, as compared to the vector-expressing clones (Fig 3C, top and bottom panels). These results are consistent with the idea that αSyn stimulates LDHA activity to shunt energy metabolism from mitochondria to glycolysis, like the Warburg Effect [29], to enable cells to cope with an energy drop once a decrease in mitochondria activity occurs.

### αSyn–expressing clones exhibit higher mitochondrial respiration and reduced mitochondrial-reactive oxygen species

Based on the results described above, we expected to find that αSyn-expressing clones will show lower levels of mitochondrial respiration. However, surprisingly, seahorse analysis showed that all αSyn-expressing clones also possess higher levels of basal and maximal respiration (OCR) as compared to the vector-expressing clones (Fig 4A, top and bottom panels). These results, showing an increase in both ECAR and OCR, are consistent with the idea that under steady state conditions αSyn acts to upregulate both energy-producing pathways converting cells to be more “energetic” (Supp Fig 1D).

**Figure 4:**
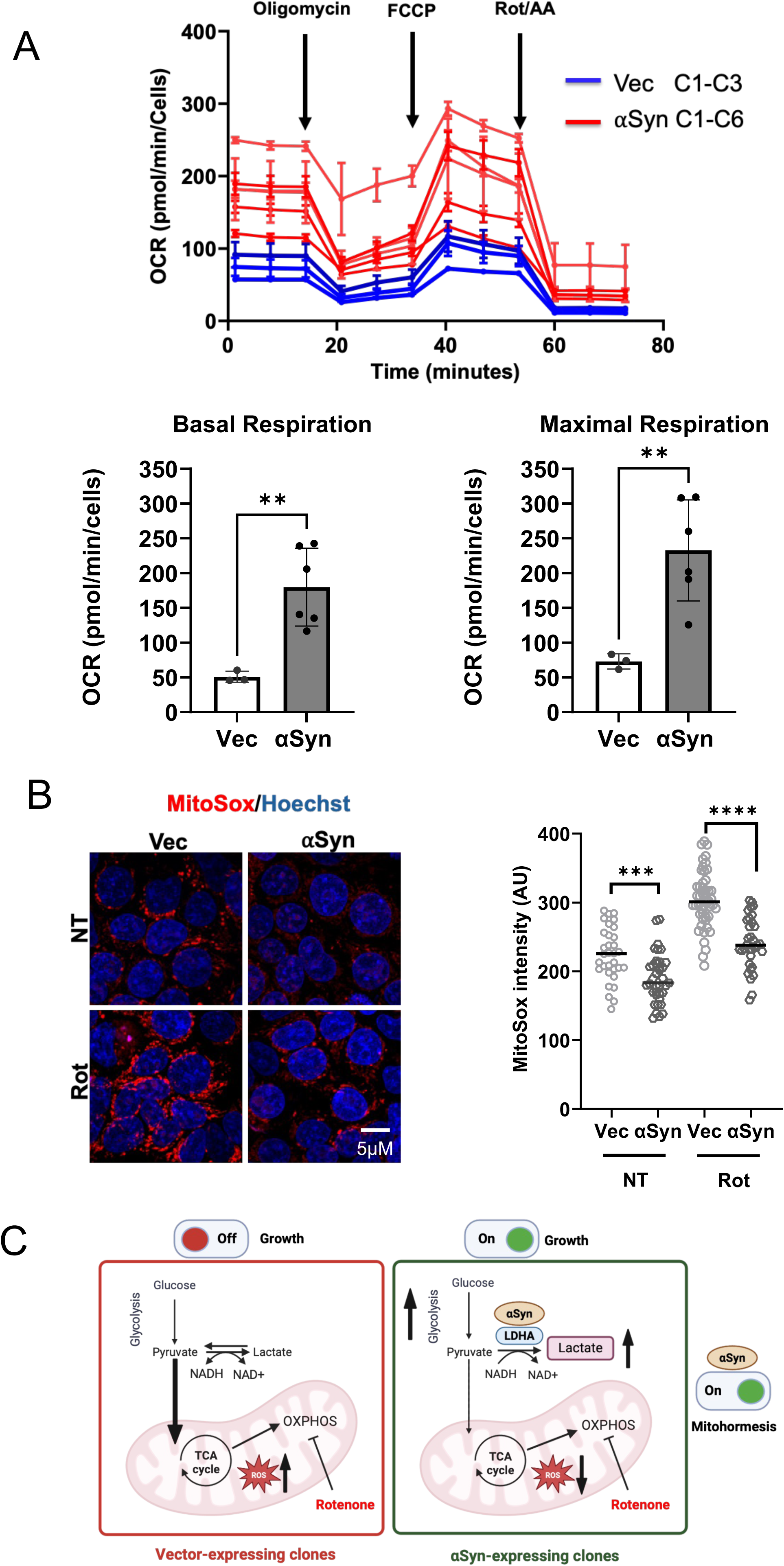
αSyn–expressing clones exhibit higher mitochondrial respiration and reduced mitochondrial-reactive oxygen species. (A) Seahorse OCR analysis of vector- and αSyn-expressing clones under basal conditions and following sequential addition of oligomycin, FCCP, and rotenone/antimycin A (AA). All αSyn-expressing clones exhibit higher basal respiration and higher maximal respiratory capacity compared to vector-expressing clones. Statistical significance: **P<0.01. (B) MitoSOX-Red analysis of mitochondrial ROS (mitoROS) in vector- and αSynexpressing clones without (NT) and with 500 nM rotenone treatment (Rot). Live cells were labeled with MitoSOX-Red (shown in red) and Hoechst (shown in blue), were imaged by confocal microscopy and analyzed using ImageJ for fluorescence intensity. *Left:* representative images. *Right:* quantification of MitoSOX fluorescence intensities from randomly selected cells in each group. αSyn-expressing clones showed lower mitoROS levels as compared to vector-expressing clones under both basal (NT) and rotenonetreated conditions (Rot). Statistical significance: ***P<0.005, ****P<0.001. (C) Schematic illustration summarizing the proposed metabolic effects in αSyn-expressing clones as compared to vector-expressing clones post rotenone treatment. *Left:* in the absence of αSyn, pyruvate is mainly routed into the mitochondria to feed mitochondrial respiration/OXPHOS. In the presence of rotenone, vector-expressing cells increase mitoROS and are “energetically cornered” resulting in their growth arrest. *Right:* In αSynexpressing clones, αSyn interacts with LDHA and stimulates its activity (increase in NAD^+^), leading to increased lactate secretion, enhanced glycolysis, and reduced mitoROS. αSyn stimulation of the LDHA-Glycolysis axis enables cells robust resistance to rotenone-induced complex I inhibition and continued growth, revealing a potent αSyn– dependent mitohormetic response.

Inhibition of respiratory complex I by rotenone results in an increase in the levels of mitochondrial reactive oxygen species (mitoROS) [4]. Thus, next we monitored the levels of mitoROS in both αSyn- and vector-expressing clones, prior and post rotenone treatment. Here we challenged cells with a high dose of rotenone to assess the protective effect of αSyn. Strikingly, MitoSox-Red quantification shows that αSyn-expressing clones possess lower levels of mitoROS, as compared to the vector-expressing clones, both prior and post high-dose-rotenone treatment (Fig 4B, left and right panels).

Thus, αSyn–expressing clones exhibit higher mitochondrial respiration and reduced mitochondrial-reactive oxygen species, coupling metabolic rewiring to improved mitohormeostasis.

## Discussion

Our findings reveal that αSyn promotes a coordinated metabolic adaptation that enhances cellular resilience to mitochondrial stress. We identify LDHA as a direct αSyn-interacting partner and show that αSyn stimulates LDHA activity, increases lactate production/secretion, augments glycolysis, reduces mitoROS, and ultimately protects cells from respiratory complex I inhibition. Collectively, these results support a model in which physiological αSyn functions as a conditional metabolic modulator that enables mitohormetic protection (see Schematic Ilustration in Fig 4C).

A central component of this model is the αSyn–LDHA–glycolysis axis. LDHA plays a key role in sustaining glycolysis, particularly during mitochondrial inhibition or hypoxia, by converting pyruvate to lactate and regenerating NAD⁺ [30, 31]. This LDHA-driven shunting of pyruvate away from mitochondria relieves metabolic overload and attenuates electron-leak-derived ROS. Our data show that αSyn enhances LDHA activity under basal conditions, positioning αSyn as an upstream regulator capable of activating a glycolytic rescue program even before overt mitochondrial dysfunction occurs.

This framework strongly parallels the well-established hypoxia–HIF-1α response, in which HIF-1α stabilization induces LDHA transcription and metabolic rewiring toward glycolysis [27]. Importantly, hypoxia was recently shown to ameliorate neurodegeneration and motor deficits in a mouse model of αSyn aggregation, despite persistence of αSyn inclusions [10]. The fact that hypoxia protects even in the presence of αSyn aggregates suggests that metabolic state, not aggregation alone, determines neuronal vulnerability. Our results raise the intriguing possibility that αSyn itself may partially mimic HIF-1αdependent metabolic remodeling, acting as an intrinsic “hypoxia-like switch” that promotes glycolytic adaptation independently of oxygen levels (Fig 4C).

The metabolic resilience conferred by αSyn is further reflected in the mitohormetic response uncovered here. Chronic low-dose rotenone exposure selectively enriched highαSyn–expressing cells and enabled αSyn-expressing clones to resist subsequent mitochondrial inhibition, whereas vector-expressing clones remained sensitive. This demonstrates that αSyn expression confers not only acute protection but also adaptive metabolic memory, a hallmark of mitohormesis [32]. The convergence between αSyn-dependent mitohormesis and classical hypoxic preconditioning suggests a shared underlying principle: adaptive metabolic rewiring that buffers cells against sustained mitochondrial stress.

Our observation that αSyn-expressing clones exhibit increased mitochondrial respiration (OCR) alongside elevated glycolysis (ECAR) may appear counterintuitive, given LDHA’s canonical role in diverting pyruvate from mitochondria. However, this dual upregulation aligns with emerging concepts in metabolic flexibility. Enhanced LDHA activity can lower NADH levels and reduce electron pressure on complex I [33], thereby suppressing mitoROS and improving mitochondrial homeostasis. At the same time, lactate produced by αSyn-expressing cells may be taken up and oxidized by neighboring cells via LDHB, that converts lactate to pyruvate, consistent with known lactate shuttling mechanisms between glia and neurons [34]. Thus, αSyn may enhance overall energetic capacity by simultaneously boosting cytosolic and mitochondrial ATP production across interacting cellular networks.

These findings carry relevance for DA neurons, which rely heavily on mitochondrial OXPHOS and exhibit high metabolic demand [35]. DA neurons are vulnerable to mitochondrial inhibition and oxidative stress, and LDHB deficiency is associated with mitochondrial dysfunction–mediated neurodegeneration [36]. In this context, αSyn-mediated activation of LDHA may serve as a protective compensatory mechanism, partially offsetting the energetic deficits caused by impaired OXPHOS. Our results showing lower mitoROS levels in αSyn-expressing clones both before and after highdose-rotenone exposure further support this model and suggest that αSyn dampens mitochondrial oxidative pressure, reducing the likelihood of ROS-driven damage.

Our work aligns with and extends prior studies reporting that αSyn interacts with glycolytic enzymes and modulates glycolytic flux. Together with the evidence that stimulating glycolysis, whether by PGK1 activation [37], hypoxia [10], or metabolic preconditioning [38], ameliorates PD phenotypes, our findings emphasize that αSyn’s physiological role may center on maintaining metabolic equilibrium under conditions of fluctuating mitochondrial performance. This contrasts with the traditional view of αSyn primarily as a toxic protein and suggests that loss of αSyn’s beneficial metabolic functions, rather than increased αSyn levels per se, may contribute to disease initiation.

In summary, we uncover a previously less-appreciated protective function of αSyn in supporting metabolic homeostasis. By binding and activating LDHA, αSyn enhances glycolysis, reduces mitoROS, increases mitochondrial respiratory capacity, and confers resistance to mitochondrial inhibition. These findings highlight αSyn as a conditional metabolic safeguard, particularly relevant for neurons dependent on OXPHOS [35], and provide a conceptual framework in which αSyn-dependent mitohormesis represents a key protective mechanism lost during PD progression.

## Materials and Methods

### Cell culture

HEK293T cells were maintained in Complete Media (CM), composed of Dulbecco’s Modified Eagle’s Medium (DMEM) with 4.5 g/L glucose (Gibco, Cat# 11965092) supplemented with 10% fetal bovine serum (Gibco, Cat#12657), 2 mM L-glutamine, 100 U/mL penicillin, and 100 µg/mL streptomycin at 37°C in a humidified incubator with 5% CO_2_. Cells were routinely tested and found negative for mycoplasma contamination.

### Generation of αSyn- and vector-expressing clones

Lentiviral particles encoding human αSyn in an αSyn–IRES–GFP vector or an empty IRES–GFP control vector (pCSC) were produced in HEK293T cells by co-transfection with the packaging plasmids pMDL, pRSV-Rev, and VSV-G using PolyJet (Cat. #SL100688, SignaGen Laboratories) according to the manufacturer’s instructions. Viral supernatants were collected 48–72 hrs post-transfection, passed through 0.45 μm filters, and used to transduce naïve HEK293T cells in the presence of 8 μg/mL polybrene. Transduced cells were expanded for 48–72 hrs, and GFP⁺ cells were sorted by FACS into 96-well plates to isolate single-cell-derived clones. Individual clones were expanded, and αSyn expression was validated by Western blotting and immunofluorescence. Vector-expressing clones were generated in parallel using the empty IRES–GFP construct.

### Rotenone treatments

Unless otherwise indicated, rotenone (Sigma, catalog #R8875) was prepared as a 10 mM stock in DMSO (Sigma, catalog #D2650) and stored at −20°C. For chronic low-dose exposure and preconditioning, cells were cultured in complete medium containing 40 nM rotenone for 2 weeks, with medium replacement every 2–3 days and then processed as indicated.

### DSG cross-linking in intact cells

Protein cross-linking was performed using disuccinimidyl glutarate (Pierce™ DSG; Thermo Fisher, Cat # 20593) as previously described [28]. Briefly, for whole-cell crosslinking, cells were washed once with PBS at 37°C and incubated with 1 mM DSG freshly dissolved in PBS for 30 min at 37°C with gentle rocking. The reaction was quenched with 50 mM Tris-HCl (pH 7.5) for 15 min. Cell pellets after cross-linking were subjected to lysis followed by immunoprecipitation or Western-blotting as described below.

### DSG cross-linking of the cytosolic S100 fraction (time course)

For the time-course cross-linking experiments, purified S100 fractions were adjusted to equal protein concentration in cross-linking buffer (PBS + protease inhibitors). DSG was added to a final concentration of 1 mM and reactions were incubated at 37°C for 0, 10, 15, 20, 30 min. Cross-linking was quenched by adding Tris-HCl (pH 7.5) to 50 mM, followed by SDS sample buffer addition and boiling. Samples were resolved by SDS– PAGE and Western blotting using anti-αSyn antibodies.

### αSyn-HA affinity purification and mass spectrometry

For identification of αSyn-associated proteins, HEK293T cells were transiently transfected with a C-terminal HA-tagged αSyn construct (αSyn–HA) using PolyJet transfection reagent (Cat. #SL100688, SignaGen Laboratories). Twenty-four hours later, cells were subjected to DSG cross-linking as described above and lysed by sonication in PBS + protease inhibitor cocktail. Lysates were cleared by centrifugation (17,000 × g, 10 min, 4°C, followed by 100,000 × g, 1 hr, 4°C) and the supernatants were incubated overnight at 4°C with rotation using anti-HA affinity resin (Thermo Fisher, Cat. #26181). Beads were washed thoroughly with PBS, and bound proteins were eluted with HA peptide (1.5 mg/mL, 2 hr at 4°C).

Eluted proteins were concentrated using Amicon Ultra centrifugal filters (3-kDa MWCO; Sigma, UFC8003), separated by SDS–PAGE, and visualized by InstantBlue® Coomassie protein staining (Expedeon, Cat. #ISB1L). Bands at ∼20, ∼43, ∼75, and ∼100 kDa were excised, digested with trypsin and desalted. The resulting peptides were analyzed using nanoflow liquid chromatography (nanoAcquity) coupled to high resolution, high mass accuracy mass spectrometry (Q Exactive Plus). The samples were analyzed on the instrument in discovery mode. The data was processed using Proteome Discoverer version 2.4, and searched against the human protein database as downloaded from Uniprot.org, to which a list of common lab contaminants was added. The search was done with two search algorithms: SequestHT and MS-Amanda. The following modifications were defined for the database search: Fixed modification- cysteine carbamidomethylation. Variable modifications- methionine oxidation, and protein N-terminal acetylation.

### Co-immunoprecipitation of αSyn-HA and LDHA-Flag

HEK293T cells were co-transfected with αSyn–HA and LDHA–Flag plasmids using PolyJet. Twenty-four hours later, cells were cross-linked with 1 mM DSG as above, quenched, and lysed in Lysis buffer (50 mM Tris-HCl pH 7.4, 150 mM NaCl, 0.5% Triton X-100, 1 mM EDTA). After clarification (17,000 × g, 10 min, 4°C), lysates were incubated with anti-FLAG Ab M2 Affinity Gel (Sigma, A2220) overnight at 4°C. Beads were washed with Lysis buffer, and bound proteins were eluted by 1.5 mg/mL Flag peptide (Sigma, F3290) for 2 hr at 4°C. Immunoprecipitates were analyzed by Western blotting with either anti-Flag or anti-αSyn Abs.

### Western blotting

Protein concentration was determined by Bradford assay (Bio-Rad). Equal amounts of protein (typically 20–40 µg per lane for whole-cell lysates; adjusted for fractions and IPs) were separated by SDS–PAGE and transferred onto Hybond PVDF membranes (Amersham). Membranes were fixed in 0.4% PFA, blocked with 5% non-fat dry milk in TBS-T (20 mM Tris-HCl pH 7.5, 150 mM NaCl, 0.1% Tween-20) for 1 hr at room temperature and incubated with primary antibodies overnight at 4°C. Primary antibodies included: anti-αSyn (15G7, Enzo Life Science), anti-GAPDH (D16H11, Cell Signaling Technology), anti-α-tubulin (YL1/2, Sigma), and anti-Flag (M2, Sigma). After washing, membranes were incubated with HRP-conjugated secondary antibodies (species-matched, Jackson Laboratories) for 1 hr at room temperature, developed with ECL reagent, and imaged using either X-Ray films (Fujifilm) or ChemiDoc Imaging system (BioRad).

### LDHA activity assay

LDHA activity was measured by monitoring NADH oxidation at 340 nm. Cells were harvested, washed in PBS, and lysed using Promega Luciferase Cell Culture Lysis Reagent (Cat. #E1531). Lysates were clarified by centrifugation (17,000 × g, 10 min, 4°C), and equal amounts of protein (1–5 µg) were added to reaction buffer containing 200 mM Tris-HCl (pH 7.3), 0.2 mM NADH, and 1 mM pyruvate. Absorbance at 340 nm was recorded at 30°C for 30 min at 1-min intervals using a Cytation 5 microplate reader (Agilent). The initial linear slope (ΔOD₃₄₀/min) was calculated for each sample. Data are presented as individual reaction slopes for each clone under basal or under two weeks 40 nM rotenone-treated conditions.

### Lactate secretion assay

Cells were seeded in 0.01% poly-l lysine (Sigma P4707) coated 6-well plates and grown to ∼70–80% confluence. After washing with PBS, cells were incubated in glucose-free DMEM (Gibco A1443001) with 10 mM glucose and 4 mM L-glutamine ± 40 nM rotenone for the indicated times. Conditioned media were collected after 8 hrs, cleared by centrifugation (1,000 × g, 5 min) and taken to lactate analysis. Lactate measurements were performed using Quris AI automated platform utilizing Jobst Technologies B.IV4 enzymatic sensors. Lactate oxidase catalyzes the oxidation of lactate, generating an electrical current proportional to lactate levels, measured and digitalized using Jobst Technologies SIX transmitter. Lactate values were normalized to total cellular protein content from parallel wells, measured by Bradford assay. Results are presented as normalized lactate levels in vector- versus αSyn-expressing clones.

### Seahorse XF analysis of glycolysis (ECAR) and mitochondrial respiration (OCR)

Glycolytic flux and mitochondrial respiration were assessed using a Seahorse Bioscience XF96 platform analyzer (Agilent Technologies, USA). For ECAR measurements, 20,000 cells were seeded in 0.01% poly-l lysine (Sigma P4707) coated Seahorse XF96 plates and incubated overnight. Before the assay, cells were washed and incubated in Seahorse XF base medium supplemented with 2 mM glutamine (pH 7.4) at 37°C in a non-CO₂ incubator for 1 hr. Basal ECAR was recorded, followed by sequential injections of glucose (10 mM), oligomycin (1 µM), and 2-deoxyglucose (50 mM) to determine glycolysis and maximal glycolytic capacity. For OCR measurements, cells were incubated in Seahorse XF base medium supplemented with 10 mM glucose, 2 mM glutamine, and 1 mM pyruvate. Basal OCR was measured, followed by sequential injections of oligomycin (1 µM), FCCP (1 µM), and rotenone/antimycin A (1 µM each) to derive parameters of basal respiration and maximal respiratory capacity. ECAR and OCR values were normalized to cell number per well.

### MitoSOX-Red staining and mitoROS quantification

mitoROS were assessed using MitoSOX Red (Thermo Fisher, Cat# M36008). Cells were grown on glass-bottom 24-well plates and treated ± rotenone as indicated. Cells were incubated with 2 µM MitoSOX Red in HBSS with Ca^2+^ and Mg^2+^ for 30 min at 37°C in the dark, washed twice with warm HBSS, and counterstained with Hoechst 33342 (for nuclei). Imaging was performed using a spinning disk confocal microscope (Nickon-CSU W1 equipped with Plan Apo λ 60x oil objective) with appropriate laser lines (MitoSOX, Ex 514/561 nm; nuclei, Ex 405 nm). Images were acquired with identical settings across conditions. Fluorescence intensity was quantified using ImageJ/Fiji: regions of interest were drawn around individual cells, and mean MitoSOX intensity per cell was measured from randomly selected fields. At least 50 cells per condition were analyzed in ≥3 independent experiments.

### Cell proliferation analysis

For proliferation assays, 2,000 cells per well were plated in 96-well plates in complete medium and monitored for 96 hours using an IncuCyte live-cell imaging platform (Sartorius). Cells were either cultured in basal medium or preconditioned for 2 weeks in 40 nM rotenone before replating into medium containing 40 nM rotenone. For control conditions, cells with the same pre-conditioning background (no rotenone or rotenone) were plated in basal media without rotenone. Cell confluence was recorded at regular intervals and fitted to an exponential growth model to calculate proliferation rate. Changes in proliferation rate were calculated as the difference between cells grown in basal media and cells grown in rotenone-containing media.

### Immunofluorescence

For αSyn immunostaining, cells were fixed in 4% paraformaldehyde for 30 min, washed in PBS, and permeabilized with 0.1% Triton X-100 in PBS for 10 min. After blocking in 5% FBS in PBS for 1 hr, cells were incubated with primary anti-αSyn antibody (dilution 1:100) for 2 hrs at room temperature, washed, and incubated with Cy3–conjugated secondary antibodies (Jackson Laboratories) for 1 hr at room temperature. Nuclei were stained with Hoechst. Images were acquired by confocal microscopy and GFP fluorescence was simultaneously monitored in the 488 nm channel. αSyn staining intensity correlated with GFP levels in αSyn-expressing clones.

### Statistical analysis

Data are presented as mean ± Standard Deviation unless otherwise indicated. Statistical analyses were performed using GraphPad Prism. Two-group comparisons were performed using unpaired two-tailed Student’s t test. P values < 0.05 were considered statistically significant.

## Acknowledgments

We thank the Weizmann BINA team - Prof. Irit Sagi, Dr. Hezi Sztainberg, Dr. Merav Marom, and Dr. Sharon Fireman - for continuous support in our studies, and the Weizmann - Azrieli Institute for Brain and Neural Sciences Research for support. A special thank you to Mr. Amnon Shoham for his support at the very early stages of our studies and to Prof. Philipp Selenko for insightful discussions, and comments on the manuscript. We also thank Dr. Elena Ainbinder from the Stem Cell Core and Advanced Cell Technologies Unit, LCSF, for providing the Seahorse facility to measure glycolysis and cellular respiration, and QurisAI for performing the lactate measurements. We thank Dr. Alon Savidor and Dr. Noga Kozer from The Nancy and Stephen Grand Israel National Center for Personalized Medicine (G-INCPM), for mass spectrometry analysis and assistance with live cell imaging and growth analysis. We are grateful to all the members of the Gross lab for their continuous support and insightful discussions.

---

**Table 1**

**LC–MS/MS analysis of αSyn–HA pulldown samples**

Table 1 summarizes the LC–MS/MS results obtained from the αSyn–HA pulldown experiment. Individual columns correspond to data derived from separate samples analyzed using two search engines (Mascot and SequestHT). Columns A–G provide aggregated annotations for each identified protein:

(A) Accession:

Database accession number for each protein entry.

(B) Description:

Protein name and annotation as listed in the sequence database.

(C) Coverage (%):

Percentage of the protein’s theoretical amino acid sequence represented by identified peptides.

(D) Proteins (#):

Number of protein entries sharing peptide evidence with the listed protein.

(E) Unique Peptides (#):

Number of peptides uniquely assigned to the protein.

(F) Peptides (#):

Total number of peptides mapped to the protein, including both unique and shared peptides.

(G) PSMs (Peptide Spectrum Matches):

Total number of MS/MS spectra matched to peptides belonging to the protein. PSMs reflect the frequency with which peptides from a protein were selected and identified during LC–MS/MS acquisition.

**Supplementary Figure 1.**
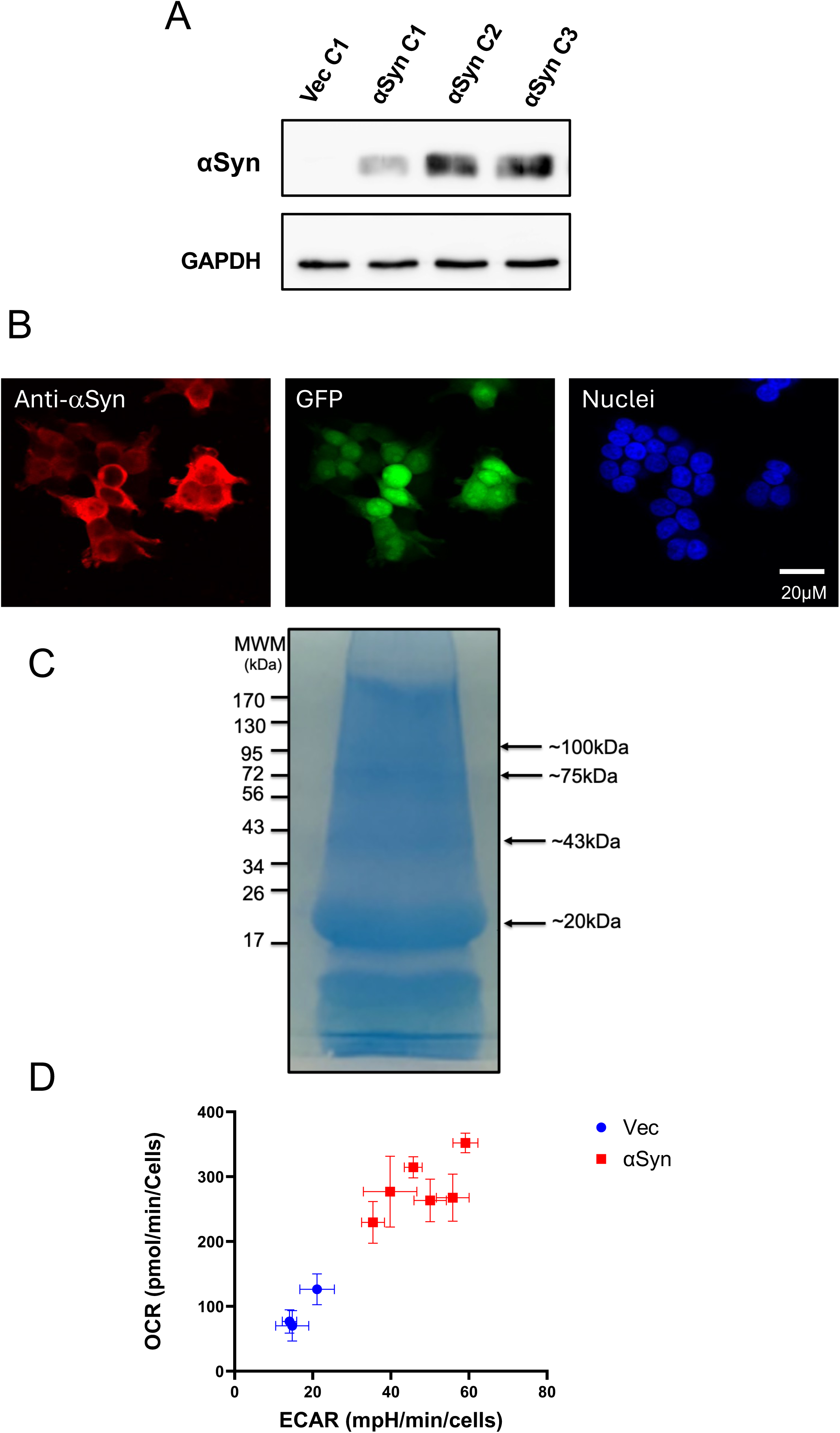
Confirmation of αSyn expression in stable clones and sample preparation for Mass Spec analysis. (A) Western blot analysis of αSyn expression with anti-αSyn Abs of one representative vector-expressing clone (Vec C1) and three αSyn-expressing clones (αSyn C1-C3) – one low expressing and two high expressing clones. GAPDH served as a loading control. (B) Immunofluorescence analysis of a representative αSyn-expressing clone. Cells were labeled with anti-αSyn antibody (red) and counterstained with a nuclear dye (blue). GFP levels were monitored in the 488 nm channel. αSyn-expressing cells display strong cytosolic αSyn staining that correlated with GFP intensity. (C) HA-tagged αSyn was transiently expressed in HEK293T cells, cross-linked with DSG, immunopurified using anti-HA antibody, eluted with HA peptide, and resolved by SDS– PAGE followed by InstantBlue® Coomassie protein staining. The indicated bands were excised and subjected to mass spectrometry analysis to identify αSyn-associated proteins. (D) Energy phenotype profile of vector- and αSyn-expressing clones. αSyn upregulates both glycolysis (ECAR) and mitochondrial respiration (OCR), shifting cells toward a more energetically active state under basal conditions.

